# Isw2-mediated chromatin remodeling governs antifungal tolerance and heteroresistance in *Candida albicans*

**DOI:** 10.64898/2026.04.21.719916

**Authors:** Maria Juliana Mantilla, Faïza Tebbji, Antony T Vincent, Louis Villeneuve, Andrew Chatr-aryamontri, Adnane Sellam

**Affiliations:** Montreal Heart Institute, Université de Montréal, Montréal, QC, Canada; Department of Microbiology, Infectious Diseases and Immunology, Faculty of Medicine, Université de Montréal, Montréal, QC, Canada; Department of Animal Sciences, Université Laval, Quebec City, QC, Canada; Institut de Recherche en Immunologie et en Cancérologie, Université de Montréal, Montréal, QC, Canada

**Author notes:** Address correspondence to Adnane Sellam.

## Abstract

Antifungal tolerance, unlike resistance, allows cells to grow slowly at concentrations above the minimum inhibitory concentration. While resistance mechanisms are well characterized, the pathways underlying tolerance remain elusive. Here, we performed a genetic screen of a transcriptional factor mutant library to identify regulators of azole tolerance in *Candida albicans*. This screen uncovered Isw2, the catalytic subunit of the ISW2 ATP-dependent chromatin remodeling complex, as a negative regulator of azole tolerance. Integrating transcriptomics, Isw2 DNA-binding profiles, and nucleosome-occupancy analyses revealed that Isw2 maintains a repressive chromatin architecture at the *CRZ1* promoter, limiting the nucleosome-depleted region and suppressing *CRZ1* transcription. Isw2 also modulates fluconazole heteroresistance and amphotericin B sensitivity through Crz1. In addition, we identified the copper-responsive transcription factor Mac1 as a context-dependent regulator of azole tolerance, acting negatively under copper limitation but positively under copper-replete conditions. Together, these findings reveal unexpected roles for chromatin remodeling and copper homeostasis in antifungal tolerance.

## Introduction

Human fungal pathogens pose a growing and often underappreciated threat to global health, causing infections that range from superficial to life-threatening systemic diseases ^1^. Collectively, fungal infections are estimated to account for 3.8 million deaths annually, with an incidence of approximately 6.5 million cases per year ^2^. In response to their increasing clinical and socioeconomic burden, the World Health Organization recently developed the Fungal Priority Pathogens List to help prioritize research and inform global public health strategies ^3^. Among the pathogens ranked in the critical priority group are several species of the *Candida* genus, notably *Candida albicans* and the multidrug-resistant *Candida auris*. These yeasts are major causes of bloodstream and invasive infections, particularly in immunocompromised individuals ^2^. Their prioritization reflects alarming trends in antifungal resistance, limited therapeutic options, and the emergence of globally distributed resistant strains, underscoring the urgent need to better understand their biology and mechanisms of drug resistance.

Therapeutic options for invasive fungal infections remain confined to three main antifungal drug classes: the polyenes, azoles, and echinocandins. Furthermore, the sustained efficacy of these antifungals is threatened by the widespread emergence of resistance, arising from heritable mutations and tolerance mechanisms that lead to treatment failure ^4,5^. Antifungal resistance arises from genetic changes or altered ploidy that allow the entire fungal population to grow at drug concentrations that would normally inhibit susceptible strains ^6^. In contrast, antifungal tolerance refers to the capacity of drug-susceptible fungal isolates to survive exposure to antifungal concentrations above the minimum inhibitory concentration (MIC), often leading to persistent infections and treatment failure ^4,6-10^. In a standard antifungal disk diffusion assay, tolerant cells appear within the inhibition zone after prolonged incubation and can be quantified by measuring the fraction of growth (FoG) ^7^. While antifungal resistance mechanisms have been extensively studied in fungi, the mechanisms underlying tolerance have only begun to be elucidated recently. Tolerance is impacted by intrinsic genetic variation among isolates of the same fungal species, as well as by environmental conditions and the tested antifungal drug. The determinants of tolerance are therefore complex, encompassing both a genetic component that varies between isolates and a non-genetic component shaped by environmental cues and cellular physiology ^6,11^. Overall, both plasma membrane and cell wall stress-response pathways, together with vacuolar trafficking and iron metabolism, play an important role in modulating fungal tolerance ^6,7,12,13^. The fungal calcium-activated calcineurin pathway plays an essential role in mediating cell wall and membrane integrity ^14-17^. In *C. albicans*, inhibition of this pathway using cyclosporine A or FK506, or genetic inactivation of the transcription factor *CRZ1*, the main effector of calcineurin signaling, completely abolished tolerance ^11^. Furthermore, pharmacological inhibition of additional pathways, including the protein kinase C (PKC) cell wall integrity signaling, the Target Of Rapamycin (TOR), the unfolded protein response (UPR), sphingolipid synthesis, Hsp90 and iron uptake, also reduced azole tolerance, underscoring the complex and interconnected regulatory network that governs fungal tolerance ^11^. Azole tolerance can also arise from alterations in ploidy and copy number variation (CNV). Recent studies have shown that both segmental and whole-chromosome aneuploidies, particularly involving Chromosome R, are associated with increased tolerance to multiple azoles ^6^. However, this form of acquired tolerance is transient, as it is lost upon the disappearance of the aneuploid chromosome or CNV ^6,18^. Although several pathways have been implicated in intrinsic fungal tolerance, the detailed mechanisms and how these pathways intersect remain largely unclear.

In the present study, we performed a genetic screen of a transcription factor mutant library to uncover novel regulators of azole tolerance in *C. albicans*. This screen identified Isw2, the catalytic subunit of the ISW2 ATP-dependent chromatin remodeling complex, as a negative modulator of azole tolerance. Integration of *isw2* mutant transcriptomic data with Isw2 genome-wide DNA-binding and targeted nucleosome-occupancy revealed that this chromatin remodeler regulates the boundaries of the nucleosome-depleted region (NDR) at the *CRZ1* promoter to mediate a closed chromatin state, thereby restricting *CRZ1* transcription. Beyond tolerance, our work delineates a role for Isw2 in modulating azole heteroresistance through Crz1. We further show that Isw2 influences amphotericin B sensitivity via Crz1, likely due to misregulation of genes encoding different vacuolar-associated functions. Our screen also identified *mac1*, a mutant lacking the copper (Cu) homeostasis regulator Mac1, which exhibited increased azole tolerance. Notably, reducing Cu levels in the growth medium markedly enhanced *mac1* tolerance, whereas Cu repletion diminished it. Together, these findings reveal a novel role for chromatin remodeling and Cu homeostasis in shaping antifungal tolerance in *C. albicans*, providing the foundation for the mechanistic analyses presented in this work.

## Results

### Genetic screen for transcriptional regulators of azole tolerance

In this study, we used the synthetic complete (SC) medium rather than the conventional rich YPD (Yeast extract-Peptone-Dextrose) to identify regulators of azole tolerance. Although the fraction of growth (FoG) is more pronounced on YPD, making it suitable for detecting mutants with reduced or absent tolerance ^11^, SC medium exhibits lower baseline tolerance ^19^ and is therefore better suited for quantitatively resolving mutants with increased tolerance. Guided by this rationale, we screened a library ^20^ of 165 transcriptional regulator mutants to identify negative modulators of azole tolerance using a disk diffusion assay and quantified tolerance based on the FoG within the inhibition zone ^7,21^. For each mutant, images of the inhibition zone were analyzed, and FoG_50_, defined as the fraction of growth within the region exhibiting 50% growth inhibition in the presence of fluconazole, was calculated using the DiskImageR software ^21^. Mutants displaying filamentous morphology were excluded from the analysis, as the increased pixel intensity of the imaged colonies reflects morphological abnormalities rather than colony growth (**Figure 1A** and **Supplementary Table S1**). Similarly, highly resistant strains were omitted because a measurable inhibition zone could not be defined for FoG_50_ calculation. This screen identified four mutants (*mac1, cta4, isw2*, and *sef2*) that exhibited significantly increased azole tolerance, two of which (*mac1* and *isw2*) were validated using independent knockout strains (**Figure 1B-C**). Loss of *ISW2*, which encodes the catalytic subunit of an ATP-dependent chromatin remodeling complex, markedly increased FoG_50_ in both SC and YPD, underscoring Isw2 as a negative regulator of azole tolerance in *C. albicans* (**Figure 1B-C**). Moreover, deletion of *MAC1*, a transcription factor governing copper uptake in *C. albicans* and other fungi ^22,23^, produced an even greater elevation in FoG50 than that observed for *isw2* (**Figure 1B**). Collectively, these findings identify Isw2 and Mac1 as potent negative regulators of azole tolerance in *C. albicans*.

**Figure 1.**
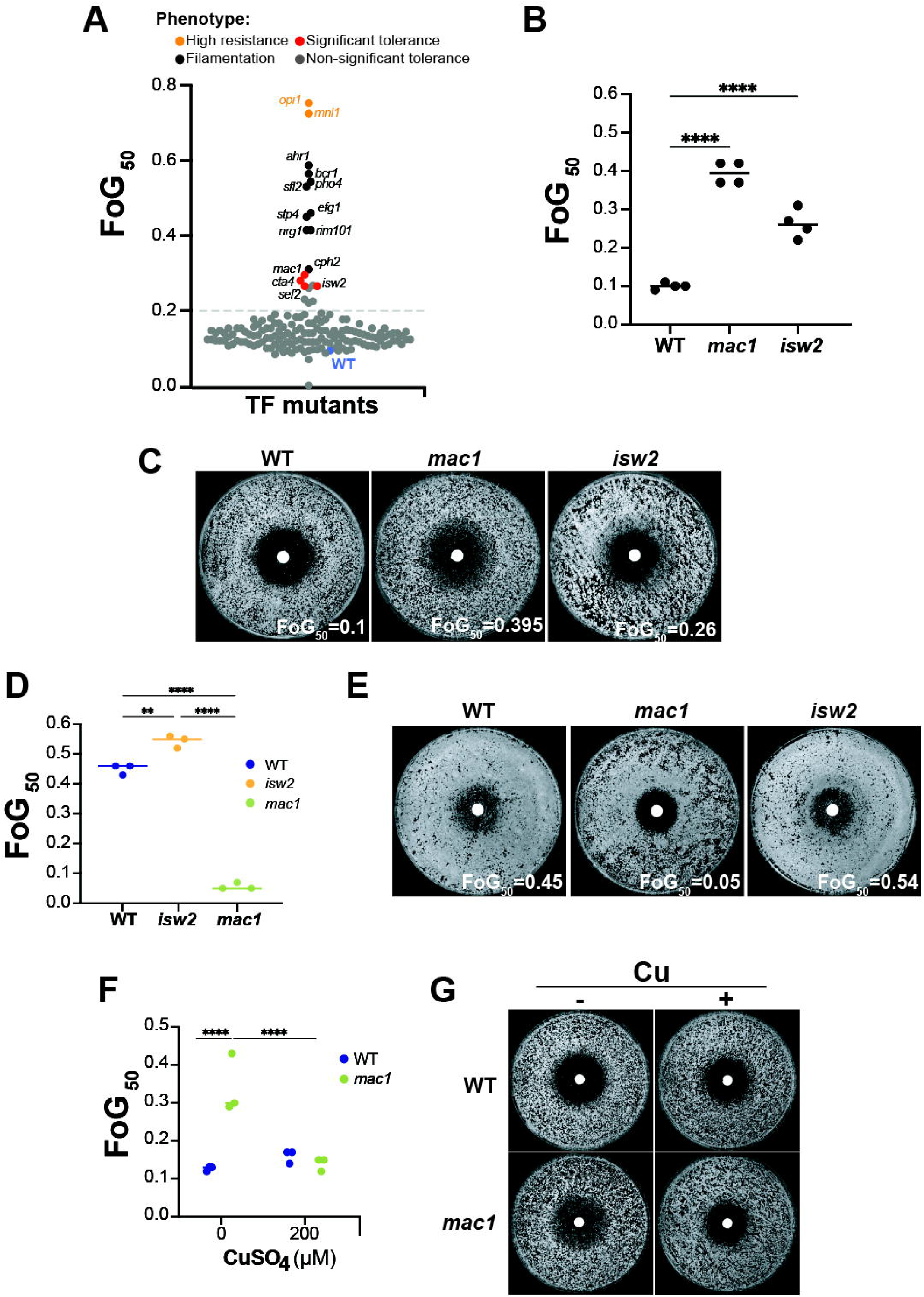
Genetic survey for transcriptional regulators modulating fluconazole tolerance. (**A**) FoG_50_ assessment of the transcription factor mutant library. For each mutant, the FoG_50_, representing the fraction of growth within the zone exhibiting 50% inhibition, was determined using the diskImageR software. (**B-C**) Validation of the tolerance phenotype of *isw2* and *mac1* mutants using SC-agar growth medium. (**B**) FoG_50_ values of WT (SN250), *mac1* and *isw2* strains grown in SC. Data represent the mean of four biological replicates. (**C**) Representative images of the disk diffusion assay for WT (SN250), *mac1* and *isw2*. Strains were assayed on SC-agar plates with disks containing fluconazole and incubated for 48h. (**D-E**) Tolerance assessment WT (SN250) and, *mac1* and *isw2* mutants using YPD-agar medium. (**F-G**) Effect of copper supplementation on tolerance of *mac1*. FoG_50_ values were determined for WT and *mac1* strains using SC-agar medium supplemented or not with 200 µM of copper (II) sulfate (**F**). Data represent the mean of three biological replicates, and representative images of disk diffusion assays are shown (**G**).

### Copper content modulates azole tolerance in *C. albicans*

*MAC1* encodes a transcription factor that controls copper (Cu) internalization in *C. albicans* and many other fungi ^22,23^. Surprisingly, FoG assays performed in YPD revealed an opposite phenotype with a marked reduction in azole tolerance compared to SC medium and the WT strain (**Figure 1D-E**). Given that YPD contains higher Cu levels than SC ^24^, we hypothesized that the increased tolerance observed in SC reflects Cu limitation in *mac1* cells, whereas the reduced tolerance in YPD results from Cu sufficiency. To test this, we assessed azole tolerance of the *mac1* and WT strains in SC medium with or without Cu supplementation. Complementing SC with 200 µM CuSO_4_ completely abolished the elevated tolerance of the *mac1* mutant, reducing it to a level comparable to that of the WT strain (**Figure 1F-G**). As Cu is also an essential cofactor for the multicopper ferroxidase Fet3, which is required for high-affinity Fe uptake ^25,26^, and because Fe availability modulates azole tolerance ^12^, we also tested the effect of supplementing this metal in SC medium. Iron addition had no impact on the increased tolerance of *mac1*, indicating that this phenotype is specifically linked to Cu availability rather than Fe-dependent processes (**Figure S1**).

### Isw2 is a negative regulator of fungal tolerance and heteroresistance

To further validate the role of Isw2 in modulating azole tolerance, a WT copy of the *ISW2* gene was reintroduced into the *isw2* deletion mutant. The resulting complemented strain displayed an intermediate FoG_50_ value between the WT and *isw2* mutant strains, consistent with the reintroduction of a single functional gene copy (**Figure 2A-B**). The sensitivity of the *isw2* mutant was also tested toward both fluconazole and miconazole using both liquid and solid assays. The *isw2* mutant displayed no discernible alterations in azole susceptibility, underscoring its specific role in modulating antifungal tolerance rather than intrinsic resistance (**Figure 2C-E**).

**Figure 2.**
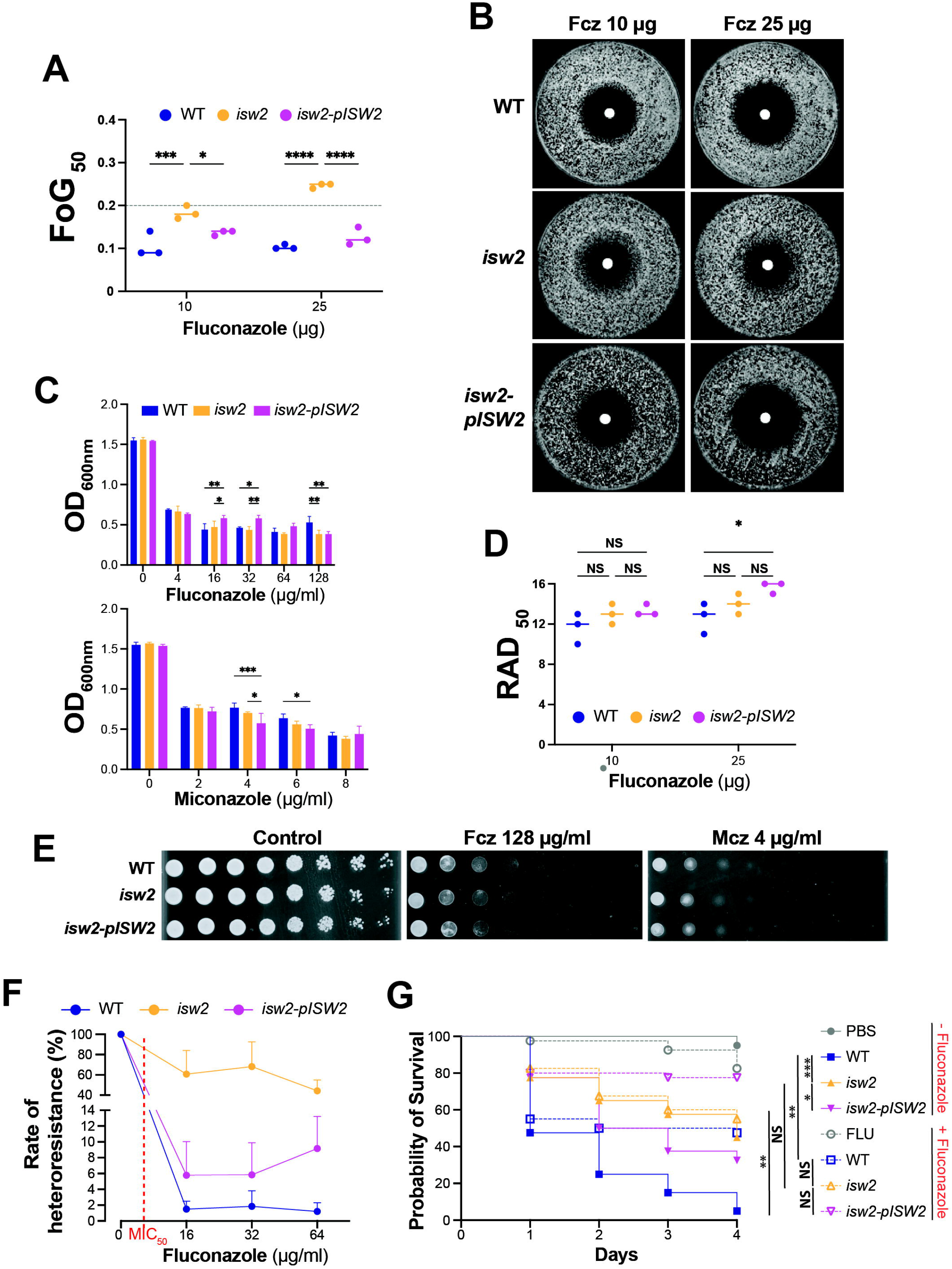
The chromatin remodeling complex ISW2 modulates tolerance and heteroresistance in *C. albicans*. (**A-B**) FoG_50_ assessment in WT (SN250), *isw2* mutant, and *isw2*-p*ISW2* complemented strain. Strains were assayed in triplicate on SC-agar plates with disks containing 10 and 25 µg of fluconazole and incubated for 48h (**A**), and representative images of the disk diffusion assays are shown (**B**). (**C-E**) Effect of *ISW2* inactivation on azole sensitivity. Effect of different concentrations of fluconazole and miconazole on WT (SN250), *isw2*, and *isw2*-p*ISW2* complemented strain was tested in liquid SC medium (**C**). Azole sensitivities were also assessed in SC solid medium using disk diffusion (**D**) and spot assays (**E**). RAD_50_ is the average radius (RAD) measured at the point where growth is inhibited by 50%. (**F**) Heteroresistance assessment using the PAP assay. (**G**) Effect of increased *isw2* tolerance and heteroresistance *in vivo* using the *Galleria* systemic infection model. Larvae were first injected with WT (SN250), *isw2* mutant, and *isw2*-p*ISW2* complemented strain, followed by a second injection with either PBS (solid line and filled symbols) or fluconazole (dashed line with open symbols). Survival curves were generated using the Kaplan-Meier method, and statistical significance was evaluated using Holm-Sidak multiple comparison test. ***P < 0.0001; **P < 0.001; *P < 0.01; NS: not significant.

Heteroresistance refers to a phenomenon in which a minor subpopulation of cells within a predominantly susceptible population displays resistance to a specific antifungal agent ^6^. Heteroresistance contributes to transient survival under drug pressure and is increasingly recognized as a key driver of antifungal tolerance, persistence, and the eventual emergence of stable resistance ^27,28^. As regulators of chromatin structure are known to contribute to cell-to-cell variability in biological systems ^29-31^, we hypothesized that Isw2 may influence azole heteroresistance in *C. albicans*. Heteroresistance of the *isw2* mutant was tested using the bacterial in-population analysis profiling (PAP) assay adapted for yeasts ^32,33^. While both WT and the *isw2*-p*ISW2* complemented strains did not grow above MIC_50_, the *isw2* mutant displayed a subpopulation of cells exhibiting a sustained growth exceeding the MIC_50_ (**Figure 2F**). Together, these data suggest that Isw2 negatively controls both azole tolerance and heteroresistance.

We further examined the *in vivo* relevance of the increased tolerance and/or heteroresistance conferred by *isw2* mutation using the *Galleria mellonella* systemic infection model. In the absence of fluconazole, larvae infected with the *isw2* mutant exhibited significantly higher survival than those infected with the WT or the complemented strain (*isw2*-p*ISW2*), indicating attenuated virulence upon *ISW2* loss (**Figure 2G**). Fluconazole treatment markedly improved survival of WT-infected larvae but produced no measurable benefit in larvae infected with the *isw2* mutant (**Figure 2G**). This lack of drug responsiveness is consistent with the intrinsic increase in azole tolerance and/or heteroresistance conferred by *ISW2* inactivation.

### Identification of the Isw2 regulon uncovers Isw2-dependent chromatin remodeling at the *CRZ1* promoter

To investigate the cellular processes related to azole tolerance that are impacted by *ISW2* deletion, we conducted transcriptomic profiling of the *isw2* mutant using RNA-seq analysis. Loss of *Isw2* resulted in extensive transcriptional deregulation, with 126 transcripts upregulated and 259 downregulated, indicating a predominant activation function for *Isw2* (**Supplementary Table S2**). This pattern contrasts with the established role of its orthologs in *S. cerevisiae* and metazoans, where they function as chromatin-based transcriptional repressors ^34-36^. Functional enrichment analysis revealed that genes upregulated in the *isw2* mutant were largely associated with metabolic processes, including nitrogen utilization, lipid biosynthesis, glycine, and heme metabolism (**Figure 3A**). Additionally, the *isw2* mutant failed to properly repress transcripts linked to oxidative stress and cellular detoxification. Genes that were downregulated were also enriched in metabolic pathways, particularly those involved in iron, sulfur, and biotin metabolism (**Figure 3B**). To determine whether these transcriptional changes reflect direct regulation by Isw2, we performed ChEC-seq to identify promoters bound by this chromatin remodeler. ChEC-seq analysis identified 1,779 Isw2 binding peaks corresponding to 894 promoter regions, representing approximately 14 % of all intergenic regions in the *C. albicans* genome (**Supplementary Table S3)**. This proportion is comparable to the genome-wide occupancy reported for Isw2 in *S. cerevisiae*, highlighting a conserved pattern of promoter targeting by this chromatin remodeler ^37^. Among these targets, Isw2 was found to associate with the promoters of 29 upregulated and 62 downregulated genes, revealing that this chromatin remodeler exerts both repressive and activating influences on gene expression in *C. albicans* (**Figure 3C**). Notably, among the direct promoter targets of Isw2, we identified the transcription factor Crz1, a well-characterized effector of the calcineurin pathway and a key regulator of azole tolerance, which was upregulated in the *isw2* mutant ^11,38^. Furthermore, many transcripts of the Calcineurin-Crz1 regulon ^15^ were significantly upregulated in the *isw2* mutant (**Figure 3D**). This observation suggests that *Isw2* may modulate antifungal tolerance by directly restricting *CRZ1* expression.

**Figure 3.**
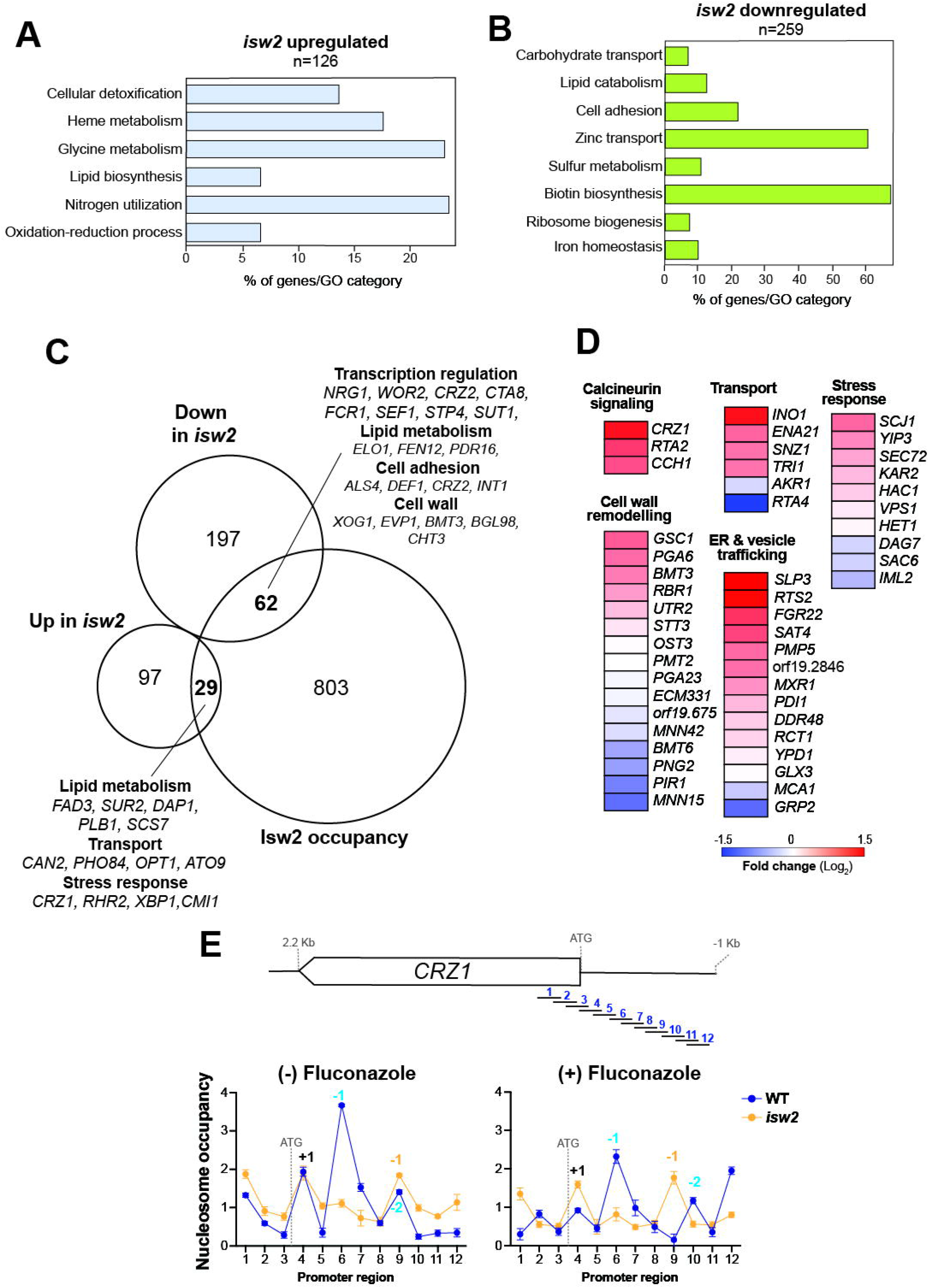
Isw2 regulates nucleosome accessibility at the *CRZ1* promoter. (**A-B**) Gene functions and biological processes enriched in *isw2* upregulated (**A**) or downregulated (**B**) transcripts. Transcripts differentially modulated were obtained by comparing the *isw2* transcriptome to that of the WT (SN250). (**C**) Isw2 regulon. Integrated analysis of RNA-seq and ChEC-seq data identifying transcripts that are differentially expressed in the *isw2* mutant and whose promoters are bound by Isw2. (**D**) Overlap of significantly upregulated transcripts in *isw2* with the set of genes defining the calcineurin regulon in *C. albicans* ^15^. (**E**) Nucleosome density defined by MNase-qPCR assay at the *CRZ1* promoter region. WT (SN250) and *isw2* cells were treated or not with fluconazole, and their DNA were digested with MNase. The position of primers spanning [-1000, +241 bp] of the CRZ1 locus is shown in the schematic. Nucleosome positions were predicted based on their occupancy peaks and their location relative to the ATG start codon. Nucleosomes differing between WT and *isw2* are shown in blue (WT) and orange (*isw2*), while nucleosomes common to both strains are shown in black. Data are presented as mean ± SD from two biological replicates, each with three technical replicates.

Since Isw2 binds the *CRZ1* promoter and governs its expression, we reasoned that this effect may be exerted through Isw2-dependent modulation of nucleosome dynamics. To test this hypothesis, we used MNase-qPCR to assess nucleosome density at the [-1000, +241 bp] region of the *CRZ1* locus in the presence or absence of fluconazole. We found that *ISW2* inactivation led to multiple alterations, most notably increased displacement of at the -1 nucleosome in both fluconazole-treated and untreated cells, suggesting an open chromatin configuration at this locus in the *isw2* mutant (**Figure 3E**). Given that the -1 nucleosome defines the nucleosome-depleted region (NDR), a structure that is dynamically remodeled to enable gene activation ^39^, our data indicate that Isw2 is necessary to regulate the NDR and promotes a closed chromatin state at the *CRZ1* promoter, thereby restricting its transcription.

### Genetic and pharmacological inactivation of the calcineurin pathway abolishes azole tolerance and heteroresistance of *isw2* mutant

To validate the involvement of the calcineurin-Crz1 regulatory axis in the enhanced azole tolerance of the *isw2* mutant, we first assessed the effect of pharmacological inhibition of calcineurin signaling using FK-506 and cyclosporin A inhibitors. The increased tolerance of the *isw2* mutant, as determined by FoG_50_, was completely abolished upon treatment with either inhibitor (**Figure 4A-B**). Similarly, deletion of *CRZ1* in the *isw2* background produced a comparable loss of tolerance. Interestingly, the heteroresistance of *isw2* was also eliminated by genetic inactivation of *CRZ1* (**Figure 4C**). Together, these findings demonstrate that Isw2 modulates azole tolerance and heteroresistance by repressing *CRZ1* transcription, thereby constraining calcineurin-dependent signaling activity.

**Figure 4.**
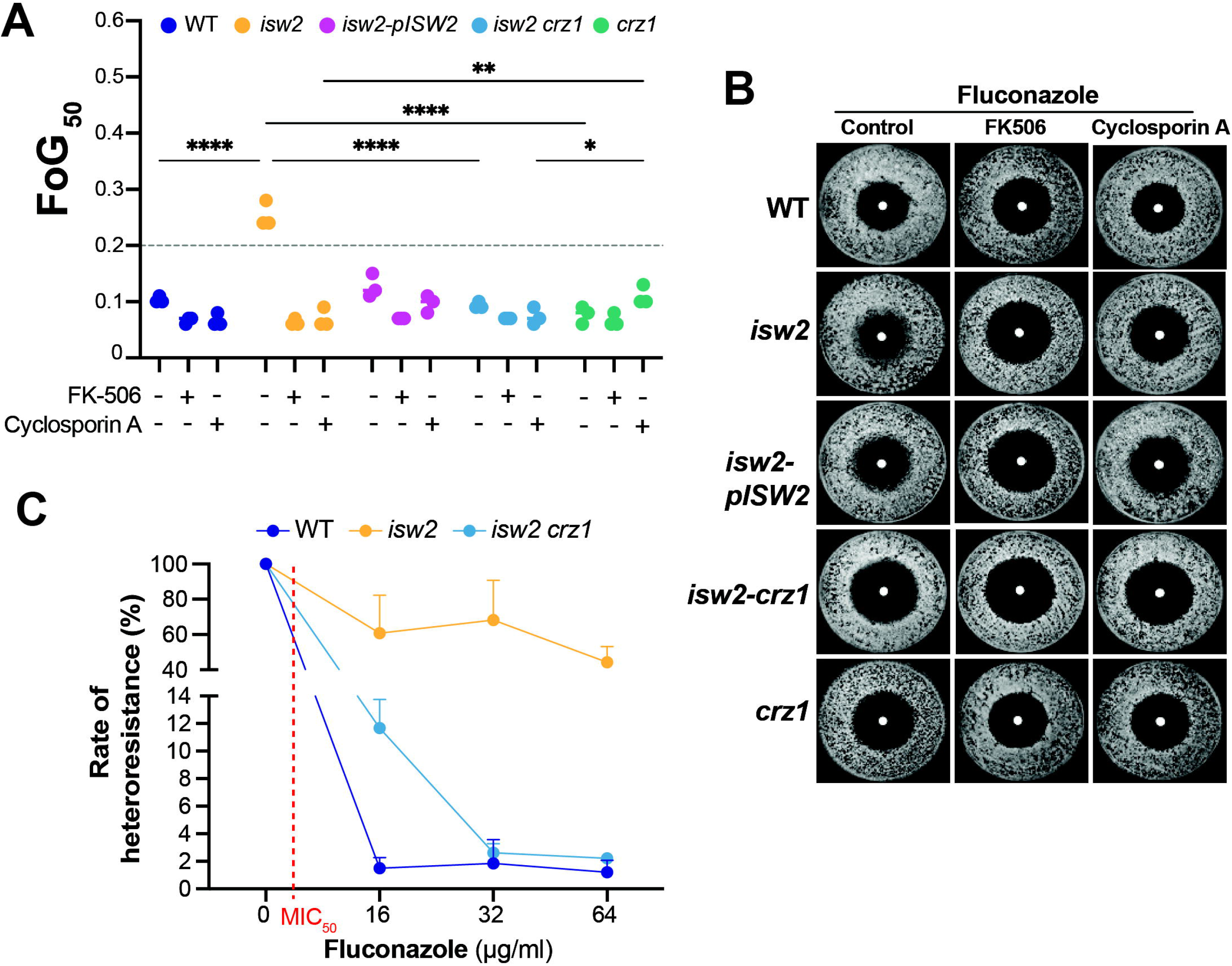
Isw2 modulates tolerance and heteroresistance through the calcineurin pathway. (**A**) FoG_50_ assessment of WT (SN250), *isw2, isw2*-p*ISW2, crz1*, and *crz1 isw2* double mutant strains in the presence of 25 µg of fluconazole. Each strain was treated or not with the calcineurin signaling inhibitors FK-506 (1 µg/ml) and cyclosporin A (1 µg/ml). (**B**) Representative images of the disk diffusion assay for strains. (**C**) Genetic inactivation of CRZ1 abolished the heteroresistance phenotype of *isw2*.

### Isw2 modulates amphotericin B sensitivity through Crz1

In addition to azole antifungals, we assessed the sensitivity of the *isw2* mutant to amphotericin B and caspofungin. While the *isw2* strain displayed WT-level sensitivity to caspofungin, it showed increased resistance to AmB (**Figure 5A-B** and **Supplementary Figure S2**). Notably, the reduced amphotericin B sensitivity of *isw2* was mediated by Crz1, as genetic inactivation of *CRZ1* restored the WT phenotype.

**Figure 5.**
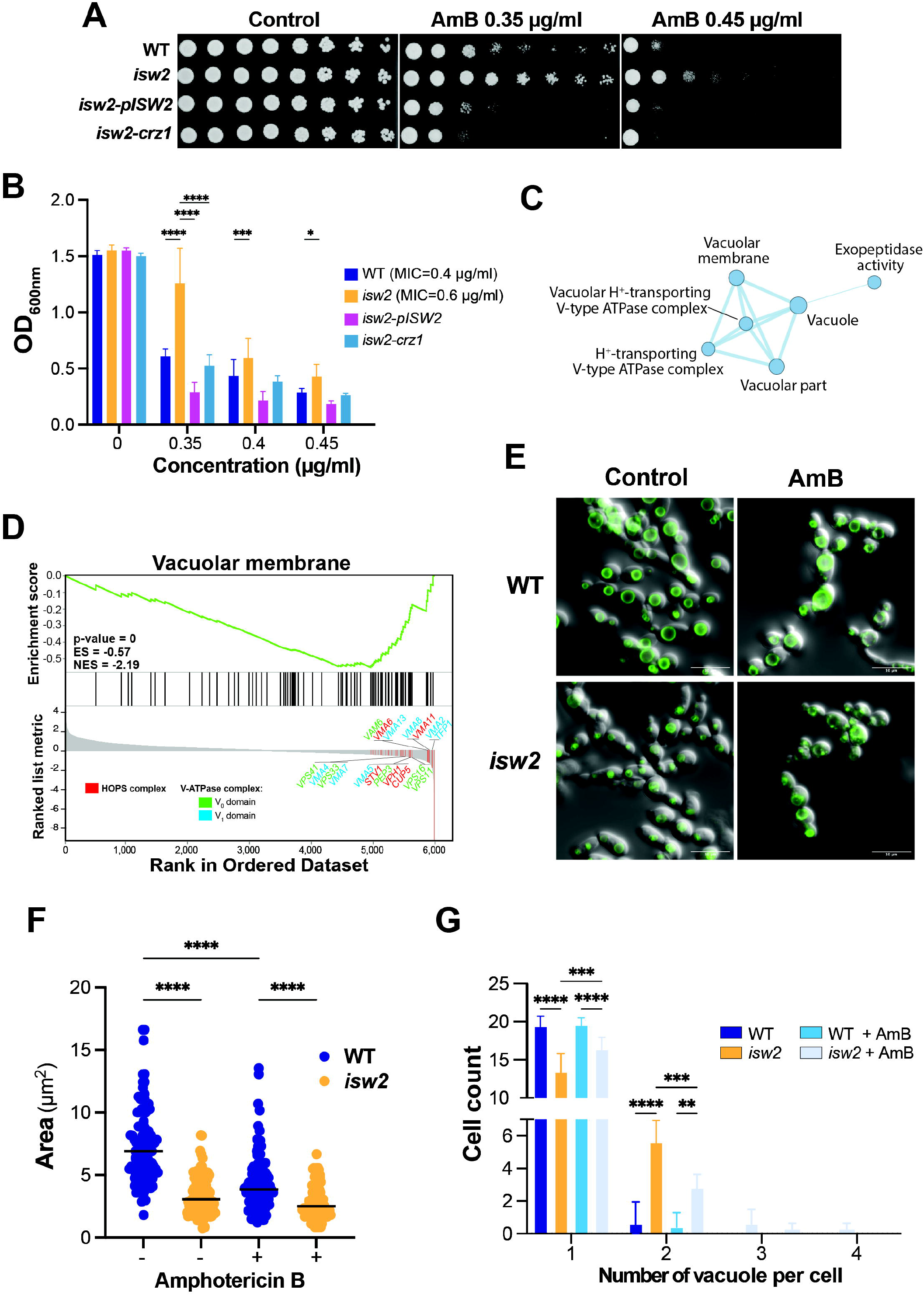
Isw2 modulates amphotericin B sensitivity. (**A-B**) The *isw2* mutant exhibits decreased sensitivity to amphotericin B. (**A**) Spot assay and OD_600_ measurements (**B**) were used to assess the effect of amphotericin B (AmB) on the growth of WT (SN250), *isw2, isw2*-p*ISW2*, and *isw2*-*crz1* double mutant. (**C**) GSEA reveals altered expression of vacuolar membrane-associated genes. The diameter of the circle reflects the number of modulated transcripts in each dataset. Images were generated using Cytoscape ^65^ with the Enrichment Map plug-in. (**D**) GSEA graphs of significant correlations between *isw2* transcriptome and vacuolar membrane gene category. Downregulated transcripts of the V-ATPase and the HOPS complexes are shown. (**E**) Altered vacuole integrity in the *isw2* mutant. *C. albicans* strains were grown in the absence or presence of 0.45 µg/ml amphotericin B and stained with the lipophilic yeast vacuolar membrane dye, FM™ 4-64. Vacuole size (**F**) and the number of each vacuole (**G**) per cell were quantified in the WT and *isw2* strains, either treated or untreated with amphotericin B.

Given that the molecular determinants underlying the sensitivity to polyene antifungals, such as amphotericin B, remain poorly defined, we investigated how *isw2* influences this phenotype by comparing the transcriptome of *isw2* cells to that of WT exposed to amphotericin B. We performed Gene Set Enrichment Analysis (GSEA) to uncover functional categories that are overrepresented among the transcripts differentially modulated in *isw2* cells. Overall, GSEA analysis revealed that transcripts involved in ribosomal biogenesis and translation were enriched in the upregulated genes, whereas genes associated with different vacuolar functions were enriched among the downregulated set (**Supplementary Table S4 and S5**). Vacuolar transcripts were enriched in processes involving the HOPS (homotypic fusion and protein sorting) tethering and the vacuolar-type proton ATPase (V-ATPase) complexes (**Figure 5C-D**). Amphotericin B is known to exert fungicidal activity in *C. albicans* and *S. cerevisiae* by targeting ergosterol in both the plasma and vacuolar membranes ^40^. To assess whether the *isw2* transcriptional pattern reflects a vacuolar defect that may underlie its reduced sensitivity to amphotericin B, we used FM4-64, a fluorescent vacuolar membrane dye, as a proxy for vacuole integrity. In WT cells, vacuoles were brightly stained along the membrane and appeared as well-defined, individualized structures (**Figure 5E**). However, in *isw2* mutant vacuoles were fragmented and exhibited a significant reduction in their size as compared the WT strain (**Figure 5E-G**). Upon treatment with amphotericin B, WT cells showed a slight reduction in vacuole size without noticeable structural changes. In contrast, *isw2* cells displayed vacuoles with similarly small and fragmented morphology as observed under untreated conditions (**Figure 5E-G**). These data suggest that the apparent amphotericin B resistance of *isw2* cells likely reflects their pre-existing vacuolar defect, which reduces the impact of the drug on the vacuole. This indicates that Isw2 modulates amphotericin B sensitivity in *C. albicans* by tuning the transcript levels of vacuolar genes and maintaining vacuole integrity.

## Discussion

In this study, we used a screening strategy biased toward identifying negative regulators of azole tolerance and uncovered previously uncharacterized roles for Isw2 and Mac1 in modulating this process. Our integrative genomic analyses demonstrated that Isw2 directly influences promoter accessibility at NDRs, thereby mediating repression of the transcription factor *CRZ1*. Furthermore, our work underscores both the similarities and the divergences in Isw2 function between *C. albicans* and *S. cerevisiae*. Isw2 occupies numerous genomic loci in both yeasts and can function as either a transcriptional activator or repressor, depending on chromatin context ^37^.

However, whereas *S. cerevisiae* Isw2 has been characterized primarily as a repressor ^34^, *C. albicans* Isw2 displays a bias toward transcriptional activation, suggesting species-specific functional adaptation of the Isw2 complex. Furthermore, as a chromatin remodeler at the promoters of the budding yeast, Isw2 was found to promote nucleosome pulling to mediate a repressive chromatin state ^41^. Consistent with this role, our data indicate that *C. albicans* Isw2 contributes to promoter chromatin organization, as loss of Isw2 resulted in an unphased -1 nucleosome. Moreover, we found that Isw2 was required to maintain +1 nucleosome occupancy specifically under drug-induced stress. Together, these findings indicate that, at the *CRZ1* promoter, Isw2-dependent modulation of −1/+1 nucleosome positioning is conserved in yeasts and supports the formation of a repressive chromatin state in *C. albicans*, particularly in response to fluconazole.

The contribution of derepressed expression of *CRZ1* to the increased tolerance of *isw2* is supported by a recent study showing that increased dosage of this transcription factor led to enhanced tolerance in *C. albicans* ^38^. Notably, this study further demonstrated that clinical isolates exhibiting increased azole tolerance display a transcriptional program consistent with constitutive activation of the calcineurin-Crz1 pathway, and that genetic loss of *CRZ1* in these isolates results in a marked reduction in azole tolerance. However, the precise mechanisms by which Crz1 and its downstream effectors promote increased tolerance remain to be elucidated in future studies. As heteroresistance rely on a small fraction of the cell population, they likely arise from cell-to-cell heterogeneity driven by differences in cell cycle stage, metabolic state, or epigenetic regulation ^42^. As a chromatin remodeler, Isw2 could interpret distinct epigenetic marks at target promoters ^43^, such as *CRZ1*, thereby promoting heteroresistance in a subset of cells.

Here we uncovered that the *isw2* mutant exhibits decreased transcript levels of vacuolar genes and, accordingly, shows an impaired structural integrity of the vacuole. Given that amphotericin B exerts its fungicidal activity through targeting ergosterol at both the plasma and vacuolar membranes ^40^, the decreased sensitivity of the *isw2* mutant to this drug is likely linked to the smaller vacuole size, which may reduce ergosterol abundance in this organelle. Consistent with this model, *C. albicans* mutants defective in ergosterol biosynthesis have been shown to exhibit reduced amphotericin B sensitivity and impaired V-ATPase function, highlighting a functional link between ergosterol and vacuolar homeostasis ^44-46^. Moreover, *erg* mutants phenocopy strains lacking individual V-ATPase subunits (*vma*^*-*^), reinforcing this relationship ^45^. Notably, *isw2* also displayed many *vma*^*-*^ phenotypes, including the altered vacuole morphology shown here, as well as impaired growth at alkaline pH, and increased sensitivity to metal ions and caffeine, as previously reported ^20^. We further observed that *isw2* mutant was significantly sensitive to the anti-arrhythmia drug, amiodarone, and showed altered growth on the non-fermentable carbon source (**Supplementary Figure S3**), both of which are *bona fide* characteristic of *vma*^*-*^ phenotypes. Together, these findings suggest that Isw2 contributes to vacuolar homeostasis, likely by maintaining vacuole size and structural integrity, which in turn influences ergosterol content at the vacuolar membrane. However, because Isw2 was not detected at promoters of vacuolar genes in our ChEC-seq analysis, its effects on vacuolar function are likely indirect. Finally, the contribution of Crz1 to decreased amphotericin B sensitivity may result from calcium leakage caused by vacuolar disruption ^47,48^, which in turn activates calcineurin-Crz1 signaling and induces a membrane stress response that helps sustain membrane integrity ^17,49,50^.

Although growth conditions are known to influence azole tolerance ^11,19^, our work identifies Cu availability as a novel determinant of this phenotype. Reduced Cu levels in the *mac1* mutant led to increased fluconazole tolerance, whereas Cu repletion attenuated this tolerance. One possible explanation for this phenotype is that *mac1* cells grown in SC medium with depleted Cu, as compared with growth in YPD, exhibit reduced ergosterol levels, the cellular target of azoles, thereby conferring increased tolerance ^51^. In support of this model, previous studies have shown that Cu starvation leads to downregulation of ergosterol biosynthetic genes and reduced ergosterol content ^22,52^. Furthermore, because Cu is critical for iron uptake ^26^, cells experiencing Cu starvation may also have reduced heme levels, an essential cofactor for the heme-containing cytochrome P450 enzyme Erg11 and the direct target of azoles ^53^, which would further contribute to decreased ergosterol levels.

## Methods

### Strains and growth conditions

The *C. albicans* strains and primers used in this study are listed in **Supplementary Table S6**. All strains were cultured and maintained at 30°C on Synthetic Complete (SC) medium containing 0.17% yeast nitrogen base without amino acids, 0.5% ammonium sulfate ((NH□)□SO□), 2% glucose, 0.2% amino acid mix, and 50 µg/ml uridine, or on Yeast extract-Peptone-Dextrose (YPD) medium containing 2% Bacto-peptone, 1% yeast extract, 2% dextrose, and 50 µg/ml uridine. For vacuolar stain Yeast extract with supplements (YES) medium was used, containing 0.5% yeast extract, 3% glucose, 0.0225% L-Adenine, 0.0225% L-Histidine, 0.0225% L-leucine, 0.0225% L-Lysine, and 0.0225% uracil.

The *isw2 crz1* double mutant was constructed by replacing the entire open reading frame (ORF) with a PCR-based disruption cassette generated from pFA plasmids, as previously described ^54,55^. For the complementation strain, the full-length *ISW2* ORF together with 1 kb of its upstream regulatory region was cloned into the pDUP3 plasmid ^56^. The resulting pDUP3-*ISW2* construct was digested using *SfiI* and integrated into the *NEUT5L* locus of the *isw2* mutant.

### Drug susceptibility tests

For the liquid assays, overnight cultures of the different *C. albicans* strains grown in SC medium were diluted to an OD_600_ of 0.1 and added into flat-bottom 96-well plates at a final volume of 200 µl per well containing antifungals. Each plate included a compound-free growth control and a cell-free negative control. Growth was monitored over 48 h at 30°C using a Sunrise™ (Tecan) absorbance microplate reader under continuous agitation. For the solid media assays, overnight cultures in SC were diluted to an OD_600_ of 4, followed by 1:50 serial dilutions. A total of 2 µl of each dilution was spotted onto SC agar plates containing the indicated antifungal and incubated for 48 h at 30°C. Images were captured using the SP-imager system (S&P Robotics, Toronto). Antifungal agents were dissolved dimethyl sulfoxide (DMSO) and included: fluconazole (128 mg/ml; Cayman Chemical, catalog no. 11594-5), miconazole (8 mg/ml; Sigma, catalog no. M3512), caspofungin (8 mg/ml; Merck, Cancidas; catalog no. 02244266) and amphotericin B (1 mg/ml; BioBasic, catalog no. AD0030P).

### Disk diffusion assay

Disk diffusion assays were performed as previously described ^21,32^ with slight modifications. Overnight cultures in SC medium were diluted in distilled water to an OD_600_ of 0.02. A total of 100 µl of the diluted culture were spread onto SC agar plates using glass beads. A 6 mm-Whatman® filter paper (1500 μm thickness) was placed at the center of each plate, and 20 µl of fluconazole (1.25 mg/ml in DMSO) was applied. Plates were incubated at 30°C and photographed after 24 and 48 h using the SP-imager system. Image analysis was performed using ImageJ (version 1.54p) and the DiskImageR pipeline in RStudio (version 2024.09.1+394) ^57^. The same methodology was applied to plates supplemented with copper (CuSO□), iron (FeCl□), and tolerance-inhibiting adjuvants. FK-506 (10 mg/ml; InvivoGen, catalog no. INH-FK5-5) and cyclosporin A (5 mg/ml; Sigma, catalog no. 30024) were dissolved in DMSO and used at a final concentration of 1 µg/ml.

### Population analysis profiling test

Heteroresistance was assessed using population analysis profiling (PAP) assay as previously described with minor modifications ^32,33^. SC agar Nunc™ OmniTrays™ plates were prepared with increasing concentrations of fluconazole (16-64 µg/ml) or without drug. Overnight cultures grown in SC medium were adjusted to a density of 10^6^ cells/ml, followed by four tenfold serial dilutions in sterile water. Then, 5 µl drops of each dilution were spotted onto the plates and incubated for 48 h at 30°C. Colonies were counted for each dilution to determine the fraction of cells capable of growing at each fluconazole concentration. Values were normalized to colony counts from antifungal-free plates.

### RNA isolation and expression profiling

Overnight cultures were grown in SC medium, diluted to an OD_600_ of 0.2 in 50 ml of fresh SC medium, and incubated at 30°C with shaking at 180 rpm to mid-log phase (OD_600_ of 0.6). Cells were treated with amphotericin B at a final concentration of 0.45 µg/ml for one hour, and untreated controls were processed in parallel. Total RNA was extracted as previously described ^58^ using acid-washed glass bead lysis in a Mini-beadbeater-24 (Biospec), followed by purification using the RNeasy kit (Qiagen). RNA-seq analysis was performed as previously described ^59^. RNA integrity was verified using an Agilent 4200 TapeStation System. RNA-seq libraries were prepared using the NEBNext Ultra II RNA Library Prep Kit (NEB) and sequenced on an Illumina NovaSeq 6000 platform at Génome Québec (Centre d’expertise et de services, Montréal). Reads were processed using fastp (v0.23.2), aligned with STAR (v2.7.9a), quantified with featureCounts (v2.0.1), and analyzed using DESeq2 (v1.40.1). Gene ontology (GO) enrichment was performed using GO Term Finder from the Candida Genome Database ^60^.

### Isw2 genome-wide occupancy by ChEC-seq

ChEC-seq (Chromatin endogenous cleavage sequencing) analysis was conducted according to the procedure described by Tebbji *et al*. (76). Isw2 was first MNase-tagged using a PCR cassette generated from the pFA-MNase-ARG4 plasmid. Overnight cultures of *C. albicans* MNase-tagged and-free MNase control strains (**Supplementary Table S6**) were diluted to an OD_600_ of 0.1 in 50 ml of fresh YPD medium and grown at 30°C to log phase (OD_600_ of 0.8). Cells were harvested and washed with buffer A (15□mM Tris [pH 7.5], 80□mM KCl, 0.1□mM EGTA, 0.2□mM spermine, 0.5□mM spermidine, one tablet Roche cOmplete EDTA-free mini protease inhibitors, 1□mM phenylmethylsulfonyl fluoride [PMSF]). Cells were resuspended in Buffer A containing 0.1% digitonin and incubated for 10 min at 30°C with shaking. MNase digestion was initiated by adding CaCl_2_ to 5□mM and incubating the samples for 5 min at 30°C. Digestion was quenched by adding 250□mM EGTA. DNA was purified using MasterPure yeast DNA purification kit (Epicentre) according to the manufacturer’s instructions and resuspended in 50□μl of 10□mM Tris-HCl buffer pH 8.0. RNA was removed by adding 10 µg of RNAse A and incubating at 37°C for 20□min. Library preparation, sequencing, and peak calling were performed as previously described ^61^. The ChEC-seq raw data are available in the GEO database with the accession number GSE314360.

### Micrococcal Nuclease (MNase)-qPCR

For MNase digestion coupled with quantitative PCR (MNase-qPCR), cells were treated with fluconazole at a final concentration of 64 µg/ml for 5 min or left untreated. Crosslinking was carried out for 30 min with 2% formaldehyde at room temperature. Samples were then washed with cold water, resuspended in 1 m of freshly prepared spheroplast buffer (1 M sorbitol, 50 mM Tris-HCl pH 7.5, 5 mM β-mercaptoethanol, and 2 mg/ml zymolyase), and incubated at 37°C for 60 min with shaking. Spheroplasts were pelleted and resuspended in 200 µl of MNase digestion buffer (1 M sorbitol, 50 mM NaCl, 10 mM Tris-HCl pH 7.5, 5 mM MgCl□, 1 mM CaCl□, 0.075% NP-40, 0.5 mM spermidine, 1 mM β-mercaptoethanol) supplemented with 60U of MNase (Worthington Biochemical) and 3 µl ExoIII (10 U/µl; New England Biolabs). Digestion was performed at 37°C for 30 min and stopped with 12.5 µl STOP buffer (50 mM EDTA and 50 mM EGTA). Samples were treated with 3 µl RNase A (10 mg/ml) for 30 min at 37°C, followed by the addition of 12.5 µl 10% SDS and 10 µl Proteinase K (20 mg/ml) and incubated overnight at 65°C. DNA was purified using phenol-chloroform-isoamyl alcohol extraction ^62^. qPCR was performed using the Luna® Universal qPCR Master Mix (New England Biolabs) on a QuantStudio□3 Real-Time PCR System (ThermoFisher) following the manufacturer’s instructions. Each sample was normalized over the undigested DNA and the actin locus. Primers sequences are listed in **Supplementary Table S6**.

### *Galleria* infection assay

Virulence assays were performed using *G. mellonella* larvae obtained from Elevages Lisard (Saint-Bruno-de-Montarville, QC, Canada), as previously described ^63^. For each experiment, larvae were randomly selected in groups of 20, and those showing signs of melanisation were excluded. Overnight cultures of *C. albicans* strains grown in SC medium were washed with PBS, diluted to 10^5^ cells in 10 µl, and injected to larvae in the last left pro-leg. A second injection of either fluconazole (5 mg/kg) or sterile PBS was administered into the last right pro-leg 2 h post-infection. Infected larvae were incubated at 37°C for four days, and mortality was recorded daily. Two biological replicates of 20 larvae each were performed for survival analysis.

### Vacuole Staining

Vacuoles of *C. albicans* strains were stained using FM™ 4-64 (Invitrogen), a yeast vacuole membrane marker, according to the manufacturer’s protocol with minor modifications. Overnight cultures were diluted to an OD_600_ of 0.1 in fresh YES medium and incubated at 30□°C with shaking until OD_600_ of 0.5. FM™ 4-64 was added to a final concentration of 8 µM, and cells were incubated for 45 min at 30°C in a water bath in the dark. *C. albicans* cells were then washed with YES medium to remove excess dye, resuspended in 5 ml of the same medium, and incubated for an additional 90 min at 30 °C with shaking. In treated samples, amphotericin B was introduced 30 min before the end of the incubation period at a final concentration of 0.45 µg/ml. Cells were pelleted, resuspended in PBS, and imaged using an LSM 710 Confocal Microscope (Zeiss). Vacuole size and fragmentation were determined using ImageJ (version 1.54p) ^64^.

### Statistical analysis

All phenotyping experiments were performed with at least three biological replicates. Gene expression and chromatin-profiling assays were conducted with two biological replicates. P values < 0.05 were considered statistically significant in the graphics. A one-way analysis of variance with **** P-value < 0.00001, *** P-value < 0.0001; ** P-value < 0.001 and * P-value < 0.01 were carried out using GraphPad Prism (version 10.6.1)

## Data availability

All RNA-seq and ChEC-seq data are available in the GEO database (https://www.ncbi.nlm.nih.gov/geo/) under the accession number GSE314360.

## Acknowledgments

We thank Dr Aleeza Gerstein (University of Manitoba) and her laboratory members, Parul Sethi and Javier San Juan Galán, for their valuable assistance with the diskImageR software. This work was supported by funds from the Canadian Institutes of Health Research project grant (grant PJT-180256), the Natural Sciences and Engineering Research Council of Canada (NSERC-DG, RGPIN-2020-06375), the Canada Foundation for Innovation and the Montreal Heart Institute foundation to Dr. Adnane Sellam. Maria Juliana Mantilla was supported by a Natural Sciences and Engineering Research Council of Canada CREATE M.Sc. scholarship (EvoFunPath program). Dr. Adnane Sellam is supported by a Fonds de recherche du Québec-Santé Senior Salary Awards.

## Author contributions

MJM and AS were involved in the conception, design, analysis, interpretation of the data; they also drafted and revised the final version of the manuscript. MJM, FT and LV carried out the experiments and analyzed the results. ATV and AC analyzed the RNA-seq and the ChEC-seq data. All authors provided critical feedback and helped shape the research, analysis, and manuscript.

## Supplementary data

**Supplementary Figure S1. Effect of iron supplementation on tolerance of *mac1***. FoG_50_ values were measured for WT (SN250) and *mac1* strains in SC medium, with or without supplementation of 100□µM ferric chloride (**A**). Data represent the mean of three biological replicates, and representative images of disk diffusion assays are shown (**B**).

**Supplementary Figure S2. The *isw2* mutant exhibits a WT sensitivity to caspofungin**.

WT (SN250), *isw2* mutant and *isw2*-p*ISW2* complemented strain sensitivity toward caspofungin was tested in SC liquid medium. OD_600_ readings were taken after 48h growth at 30°C under shaking.

**Supplementary Figure S3. The *isw2* mutant displays increased sensitivity to the antiarrhythmic drug amiodarone** (**A**) **and impaired growth on the non-fermentable carbon source lactate** (**B**). (**A**) Data represent the mean growth inhibition (% relative to the untreated condition) after 24 h of growth in SC medium containing amiodarone, calculated from at least three independent replicates. (**B**) *C. albicans* cells were grown for 24 h in SC-glucose or SC-lactate medium. Data represent the mean growth percentage relative to the SC-glucose condition.

**Supplementary Table S1**. Raw data of the genetic survey for transcriptional regulators of fluconazole tolerance.

**Supplementary Table S2**. RNA-seq analysis of the *isw2* mutant. Lists of raw RNA-seq data and differentially expressed transcripts in the *isw2* mutant compared to the WT strain grown in SC medium. Differential expression was determined using a false discovery rate (FDR) of 1% and a

± 1-Log_2_ fold change cutoff.

**Supplementary Table S3**. List of Isw2-occupied loci identified by ChEC-seq.

**Supplementary Table S4**. RNA-seq analysis of the *isw2* mutant in response to amphotericin B treatment. Lists of raw RNA-seq data and differentially expressed transcripts in the *isw2* mutant compared to the WT strain grown in SC medium supplemented with 0.45 µg/ml amphotericin B for 1h. Differential expression was determined using an FDR of 1% and a ± 1-Log_2_ fold change cutoff.

**Supplementary Table S5**. GSEA analysis of the amphotericin-treated *isw2* cells.

**Supplementary Table S6**. Primer and strain lists.

