## Supplementary figures and images for "Isw2-mediated chromatin remodeling governs antifungal tolerance and heteroresistance in *Candida albicans*"

### Supplementary Figure S1

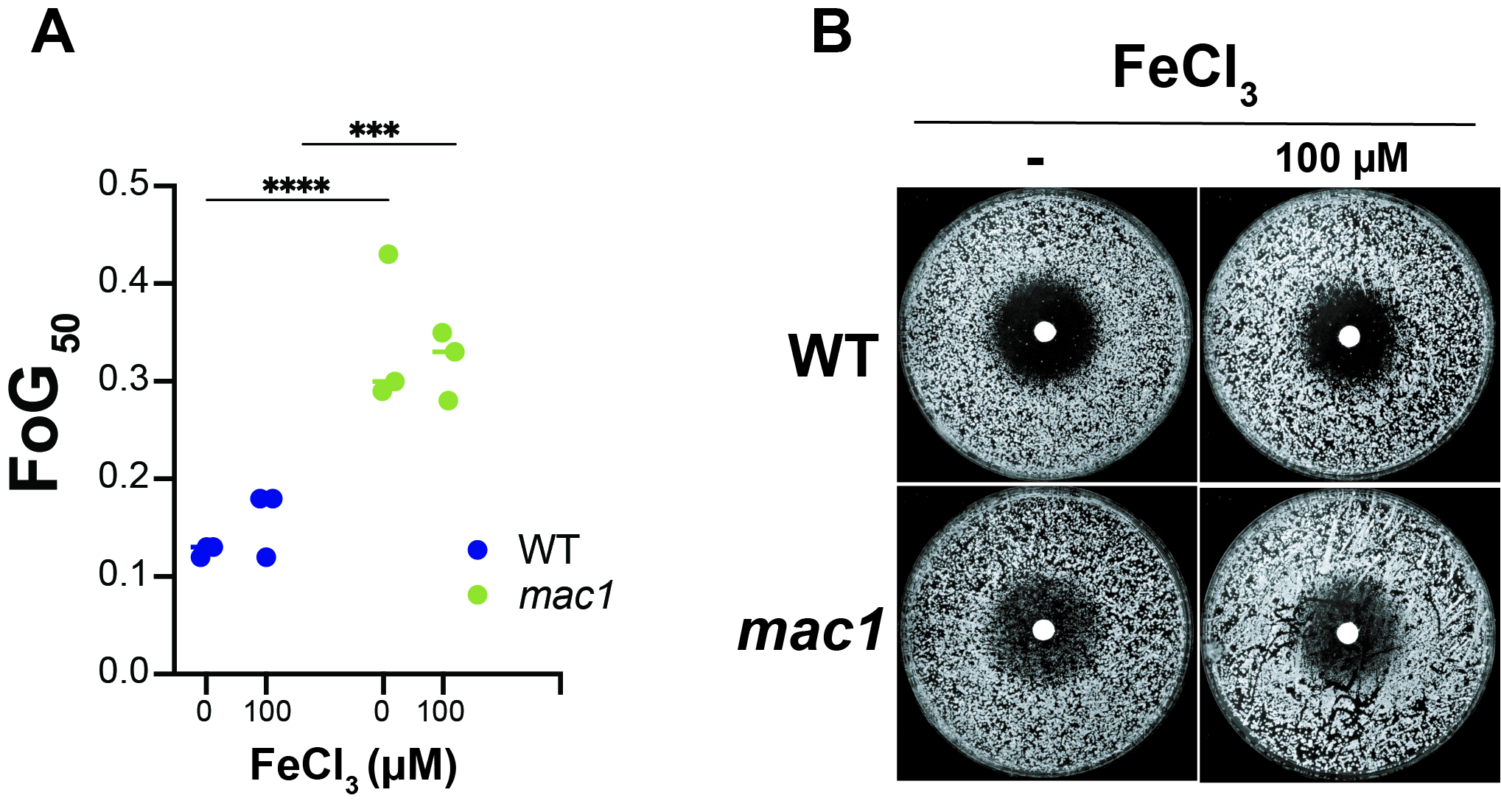

### Supplementary Figure S2

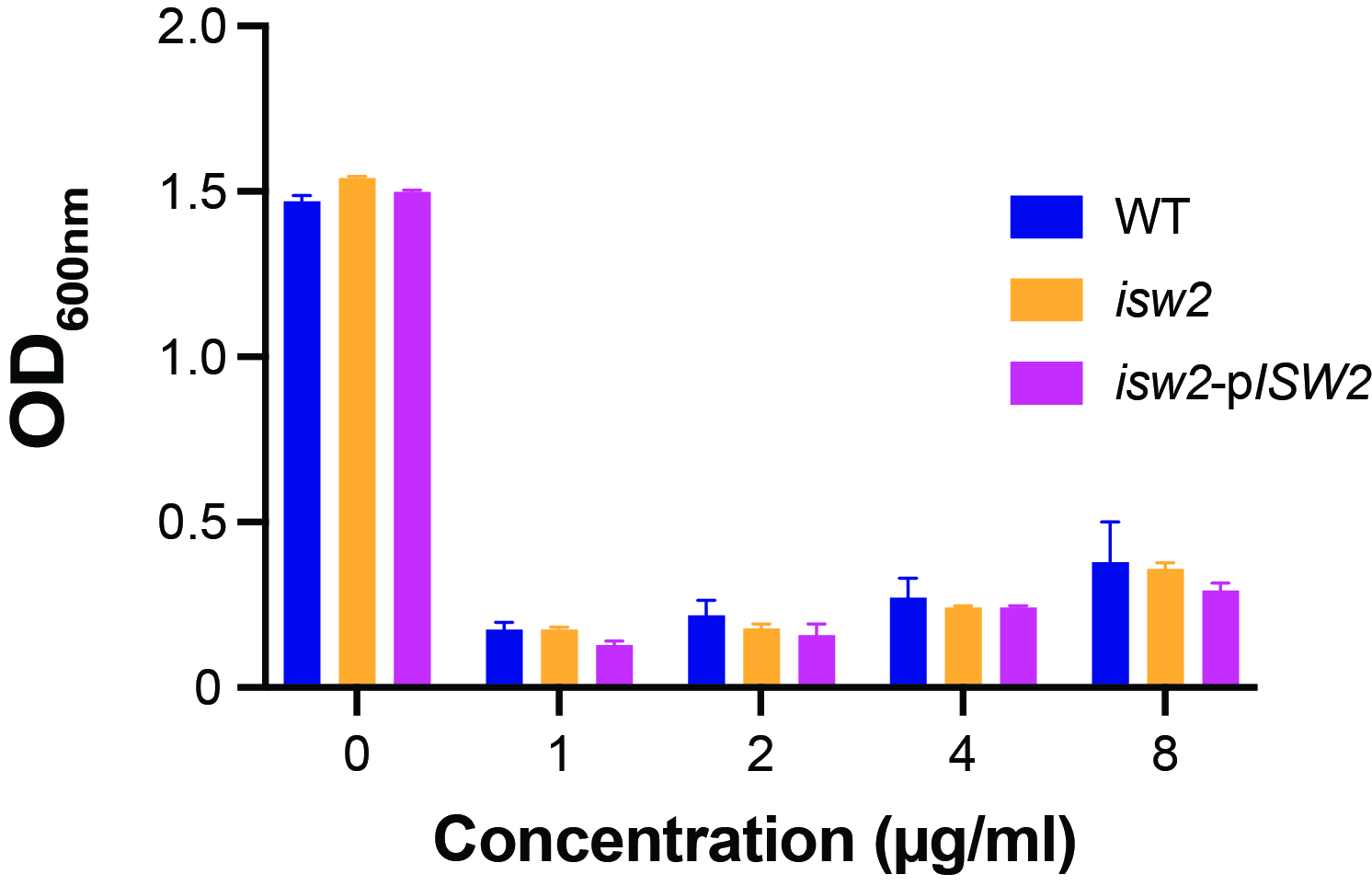

### Supplementary Figure S3

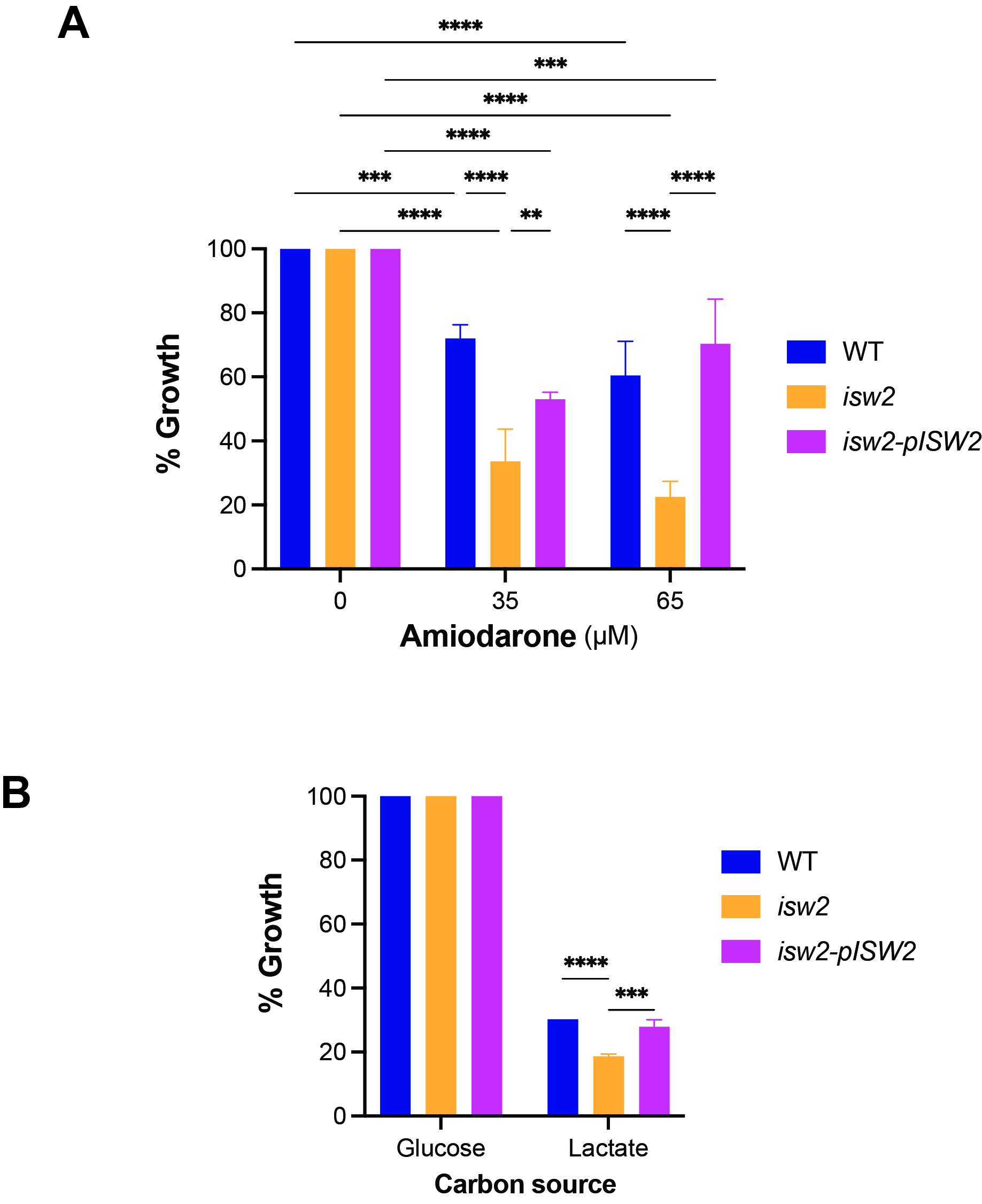
